# Model-driven characterization of functional diversity of *Pseudomonas aeruginosa* clinical isolates with broadly representative phenotypes

**DOI:** 10.1101/2023.10.08.561426

**Authors:** Mohammad Mazharul Islam, Glynis L. Kolling, Emma M. Glass, Joanna B. Goldberg, Jason A. Papin

**Affiliations:** Department of Biomedical Engineering, University of Virginia, Charlottesville, Virginia, 22903; Department of Pediatrics, Emory University, Atlanta, GA 30322

**Author notes:** Corresponding author. All correspondence and material requests should be addressed to.

**Keywords:** *Pseudomonas aeruginosa*, Phenotypic diversity, Metabolic modeling

## Abstract

*Pseudomonas aeruginosa* is a leading cause of infections in immunocompromised individuals and in healthcare settings. This study aims to understand the relationships between phenotypic diversity and the functional metabolic landscape of *P. aeruginosa* clinical isolates. To better understand the metabolic repertoire of *P. aeruginosa* in infection, we deeply profiled a representative set from a library of 971 clinical *P. aeruginosa* isolates with corresponding patient metadata and bacterial phenotypes. The genotypic clustering based on whole-genome sequencing of the isolates, multi-locus sequence types, and the phenotypic clustering generated from a multi-parametric analysis were compared to each other to assess the genotype-phenotype correlation. Genome-scale metabolic network reconstructions were developed for each isolate through amendments to an existing PA14 network reconstruction. These network reconstructions show diverse metabolic functionalities and enhance the collective *P. aeruginosa* pangenome metabolic repertoire. Characterizing this rich set of clinical *P. aeruginosa* isolates allows for a deeper understanding of the genotypic and metabolic diversity of the pathogen in a clinical setting and lays a foundation for further investigation of the metabolic landscape of this pathogen and host-associated metabolic differences during infection.

**Impact statement:** *Pseudomonas aeruginosa* is a leading cause of infections in immunocompromised individuals and in healthcare settings. The treatment of these infections is complicated by the presence of a variety of virulence mechanisms and metabolic uniqueness among clinically relevant strains. This study is an attempt to understand the relationships between isolate phenotypic diversity and the functional metabolic landscape within a representative group of *P. aeruginosa* clinical isolates. Characterizing this rich set of clinical *P. aeruginosa* isolates allows for a deeper understanding of genotypic and metabolic diversity of the pathogen in a clinical setting and lays a foundation for further investigation of the metabolic landscape of this pathogen and host-associated metabolic differences in infection.

## Introduction

*Pseudomonas aeruginosa* is a major contributor to a broad range of nosocomial infections and is commonly associated with many clinical cases including cystic fibrosis, pneumonia, urinary tract infections, sepsis, and skin infections ^1–7^. The ability of *P. aeruginosa* to survive and adapt to diverse and challenging habitats and growth conditions makes this opportunistic pathogen successful in colonizing and infecting many physiological niches within the human host ^8^. Clinically isolated *P. aeruginosa* strains are often multiclonal and demonstrate diverse metabolic and phenotypic traits. Many recent studies investigated strain-specific and condition-specific gene essentiality ^9^, antibiotic susceptibility ^10,11^, horizontal gene transfer, virulence and emergence of antimicrobial resistance ^12–16^, and phylogenetic ^17^ and phenotypic diversity ^18^ of clinically isolated strains. Additional work explored the functional repertoire of the core and accessory genomes in *P. aeruginosa* ^9,13,19–21^. These studies broadened our understanding of the genomic landscape across the *P. aeruginosa* pangenome and antimicrobial resistance in strains from a wide range of infection types and environments.

However, most of the abovementioned studies either only focused on the genetic diversity among clinical isolate populations or selected isolates from a single infection site or environment and sometimes only interrogated specific clonal types of interest. Studies that included a large population of isolates often selected a smaller subset based on SNP diversity within the isolate sequences. Therefore, comprehensive investigations of the diverse phenotypes and functional metabolic repertoire of clinically isolated strains of *P. aeruginosa* from multiple body sites and associated with multiple patient comorbidities are still rare. Moreover, genetic variability, mutations, and recombination events play an important role in the phenotypic diversification and metabolic heterogeneity of *P. aeruginosa* ^22,23^, which is also evident in the phenotypes observed in our previous study with these isolates ^10^. These limitations have made it difficult to gain a deeper understanding of *P. aeruginosa* metabolism and infections that can span many isolation sites in the human body, multiple patient comorbidities, and other host factors, that can affect the observed microbial phenotypes.

We hypothesize that the metabolic differences across *P. aeruginosa* isolates are dependent on a complex combination of host influences and pathogen-specific factors, which can be delineated using a combination of omic analyses and the experimental characterization of unique metabolic traits in different clinical isolates. The advancements in efficient, high-throughput genome sequencing have opened the possibility to better understand the diverse metabolism of *P. aeruginosa* and the relationships between multi-omics data. To this end, genome-scale metabolic modeling has been used in previous studies to explore the functional diversity among strains of a species or within a pangenome ^24,25^.

In this work, we performed experimental and computational analyses including sequencing and analyzing genomes of a set of clinical isolates representative of the diversity observed in clinical settings. We explored the genetic, sequence type, and phenotypic diversity and their correlation within the representative set of isolates. We performed functional annotation of these genomes to enable the reconstruction of genome-scale metabolic networks, and we performed flux sampling analyses with these network models to delineate differences and similarities of the *P. aeruginosa* clinical isolates. We evaluated metabolism shared across isolates as well as metabolism unique to each. By providing an integrated view of the correlation between clinical metadata information (patient demographics, isolation source, and isolate morphology), and high-throughput multi-omics data, we can better characterize the functional diversity of clinical isolates, and hopefully better understand how to treat infections caused by this versatile human pathogen.

## Results and Discussion

### Genome sequencing of the representative isolate group

Between February 2019 and February 2020, a total of 971 clinical *P. aeruginosa* isolates from 590 patients at the University of Virginia (UVA) Health system, were collected as reported previously ^10^. For each of the 971 clinical P. aeruginosa isolates, patient demographic profiles (age, sex), comorbidities (cystic fibrosis or diabetes), and isolate phenotypic traits (mucoid phenotype, metallic sheen, pigment production, and hemolytic activity) were collected and tabulated. To understand their genotypic, phenotypic, and metabolic variance and to assess the shared and unique traits that they can manifest, we extensively studied a sample population of 25 of these *P. aeruginosa* isolates. We performed stratified random sampling (see Methods section for additional details) to identify a sub-population of 25 isolates which represents the phenotypic diversity of the entire population of 971 isolates. Figure 1 shows the schematic of the stratified random sampling procedure (Figure 1A), a visual representation of their sequence similarity analyses (Figure 1 B), and the phenotypic properties and associated patient metadata of the 25 selected isolates (Figure 1 C).

**Figure 1:**
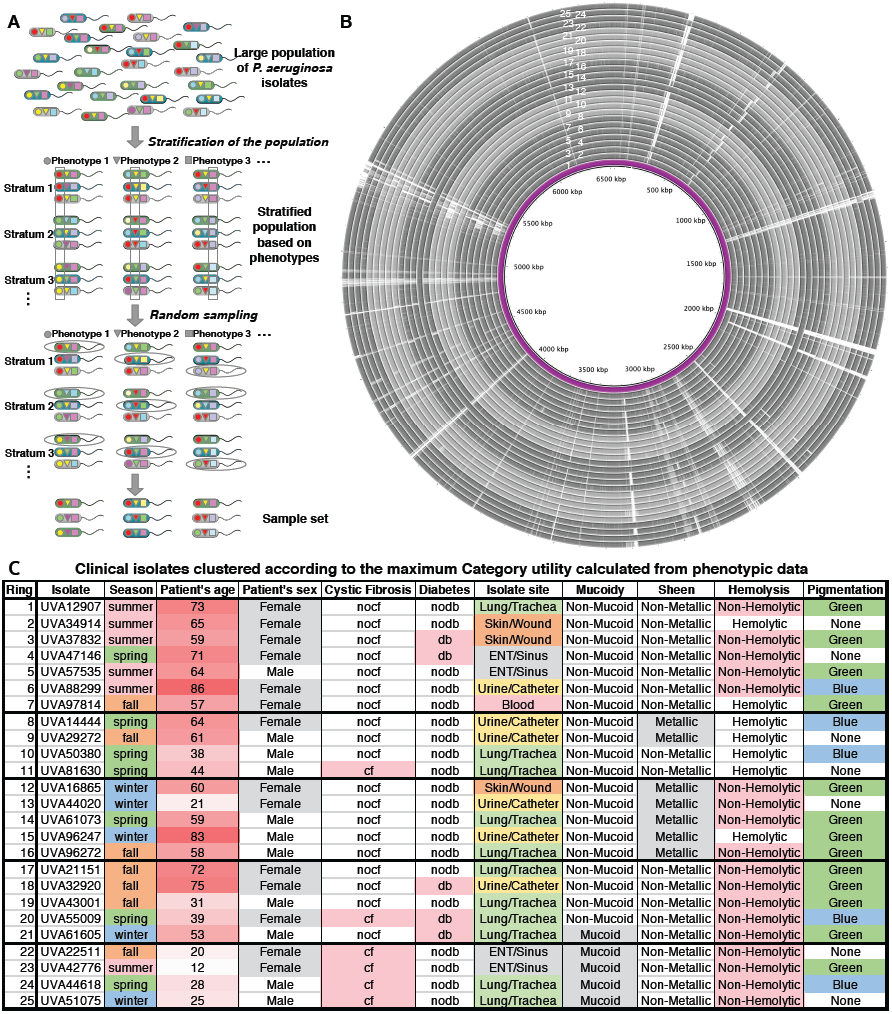
Process of selecting a representative set of 25 clinical isolates from the large population, their genome similarities, and phenotypic clustering. (A) Schematic of the stratified random sampling procedure to select a representative set of 25 isolates from the large isolate collection by Dunphy *et al*. ^10^ ^27,28^**(B)** Circular comparison of the isolate genomic sequences with PA14 genome sequence as reference (innermost circle) generated by BLAST Ring Image Generator ^29^. BLAST rings are arranged in five groups of alternating light and dark grey based on the phenotypic categories from panel C. **(C)** Morphology and patient metadata associated with the selected 25 isolates. Isolates are grouped into 5 clusters based on their phenotypic categories (category utility value = 0.3999). Phenotypes are color coded according to the category values. “*nocf*” = patient does not have Cystic Fibrosis, “*cf*” = patient has Cystic Fibrosis, “*nodb*” = patient is non-diabetic, “*db*” = patient is diabetic.

Isolates were grouped into five clusters based on their phenotypic similarities using the measure of Category Utility (see Figure 1C). The category utility hypothesis proposed by Corter and Gluck ^26^ states that the categories that become preferred in a population are those that best describe the diversity of the population (see Methods section for additional details). We used Category Utility to maximize both the probability that isolates in the same cluster have phenotypic attributes and patient metadata in common, and the probability that isolates from different clusters have different phenotypic attributes and patient metadata. This is a measure of the probability that all the isolates in each cluster will assume the same categorical value for each of the 10 different phenotypic categories in this study. The maximum calculated category utility value achieved for this clustering (in panels 1B and 1C) was 0.3999 out of 125 different random combinations of clusters. With this clustering, then, there is approximately a 40% probability that all the isolates grouped together have the exact same values in every phenotypic property, and also that isolates in different groups do not have the same values in every phenotypic property.

Figure 1A illustrates the stratified random sampling procedure for selecting the subset of clinical isolates representative of the phenotypic diversity of the entire population of 971 isolates. First, the isolate collection is separated into different strata based on each of the 10 phenotypic properties (season, patient’s age, patient’s sex, isolation site, presence of cystic fibrosis and diabetes, mucoidy, hemolytic capability, metallic sheen, and pigmentation). In each stratum, the number of isolates maintains the same proportion as the original large population. Then isolates are chosen at random from each stratum and a sample set of 25 isolates is chosen.

To get the complete genomic and functional profile of the 25 selected isolates, we sequenced their genomes. The comparison of the whole genome sequences of the 25 isolates was performed using the MEGA11 software^27^ (see Methods section for additional details). We aligned the aligned genome sequences to the reference *P. aeruginosa* PA14 genome (NCBI ID: NC_008463.1) and the alignment is displayed in a ring diagram. The purple ring in the middle of the ring diagram is the PA14 reference genome. The rings are ordered according to their phenotypic clustering shown in panel C. The gaps in the rings correspond to the gaps in the sequence alignments.

### Correlation between whole genome sequence, sequence types, and the phenotypic properties of the isolates

Since the isolates demonstrate some degree of genetic variability, their functional uniqueness might play an important role in the phenotypic diversification and metabolic heterogeneity. The wide range of phenotypic diversity observed in our previous study with these isolates ^10^ led us to investigate further the correlation between the genome sequence of the isolates and their phenotypes. In addition to the diversity based on whole-genome sequences, multi-locus sequence typing (MLST)^30^ was used to evaluate the allelic variation of seven housekeeping genes in the isolate genome sequences, generating the sequence types (STs) to characterize isolates. The sequence types of the isolates are listed in supplementary information 1.

Figure 2 A shows the hierarchical clustering of the *P. aeruginosa* clinical isolates based on their sequence types and similarity scores on the seven housekeeping genes (allelic profiles). The color scale shows that the Euclidean distances between most of the isolates are low, except isolate UVA88299 setting itself apart in the distance matrix, indicating that its allelic profile is the most distinct from the other isolates. The two other most distinct isolates are UVA51075 and UVA61605, which are also very distant from each other.

**Figure 2:**
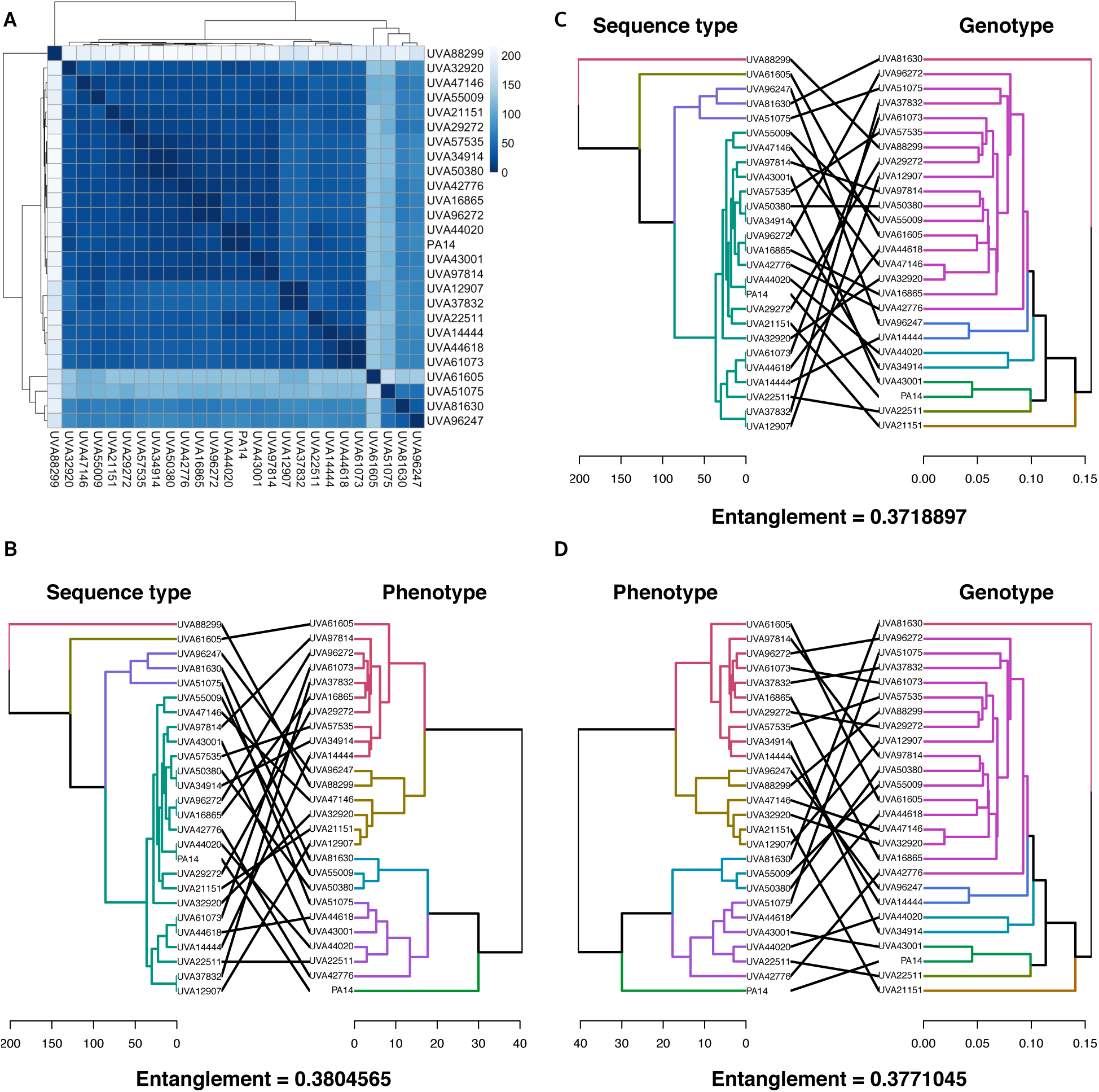
Clustering of the *P. aeruginosa* clinical isolates based on multi locus sequence types, genome sequence, and phenotypes, and their correlation with each other. (A) Hierarchical clustering of the isolates based on their multi locus sequence type (MLST) based on seven housekeeping genes. Comparison of clustering between (B) MLST and Phenotype, (C) MLST and Genotype, and (D) Phenotype and Genotype.

To investigate if the allelic profiles of the isolates are related to the genomic differences based on whole genome sequences of their phenotypic clustering, we compared the hierarchical clustering of the isolates based on the MLST (allelic profile) to the phenotypic clustering (Figure 2B) and genotypic clustering (Figure 2C). In addition, we compared the genome sequence-based clustering to the phenotypic clustering to understand their correlation (Figure 2D). These comparisons were performed by superimposing the two respective hierarchical clustering trees and enumerating the entanglement scores between them. A lower (closer to zero) entanglement score means that the trees are very similar to each other and the correlation between the two respective hierarchical clustering is good.

It is noticeable from the superimposed trees and the entanglement scores that the phenotypic, genotypic, and MLST-based clustering do not correlate well with each other. The observed poor correlation may indicate significant differences in functional annotation differences, regulatory mechanisms, post-translational modifications, and other factors that define the functional behavior of an organism in a given environment. We also note that the entanglement scores are highly similar in each of the comparisons; the phenotype to genotype similarity is comparable to the sequence type to genotype similarity which suggests that neither the genotype nor the sequence type are strong predictors of the observed phenotypes. To further characterize the diversity among the clinical *P. aeruginosa* isolates, we performed functional annotation of the isolates’ genome sequences.

### Functional annotation of the isolates and *P. aeruginosa* pan-genome analysis

Each of the isolate genome sequences was annotated for metabolic functions based on the KEGG biochemical database ^31,32^ (see Methods section for details). For each isolate, an average of 50,000 BLAST hits were found in organisms across 1462 genera, 506 families, and 61 phyla from the KEGG database. Interestingly, most of the metabolic functions in the *P. aeruginosa* isolate genomes are shared with >1000 organisms in the KEGG database, while only a few metabolic functions in the isolates are shared with <10 organisms. Figure 3A shows a distribution of KEGG orthologues (KO) annotated in the isolate genomes and the number of non-*P. aeruginosa* species the specific KO is shared with. Out of the approximately 2000 KOs in the *P. aeruginosa* isolates that were found to be shared with species in the KEGG database, only 27 were shared solely with other *Pseudomonas* species. These KOs mainly belong to the biosynthesis of macrolides, phosphonate and phosphinate metabolism, and biosynthesis of polyketide sugar units. As indicated with the axis on the right of Figure 3A, all KOs in the profiled *P. aeruginosa* isolates are found in other organisms.

**Figure 3:**
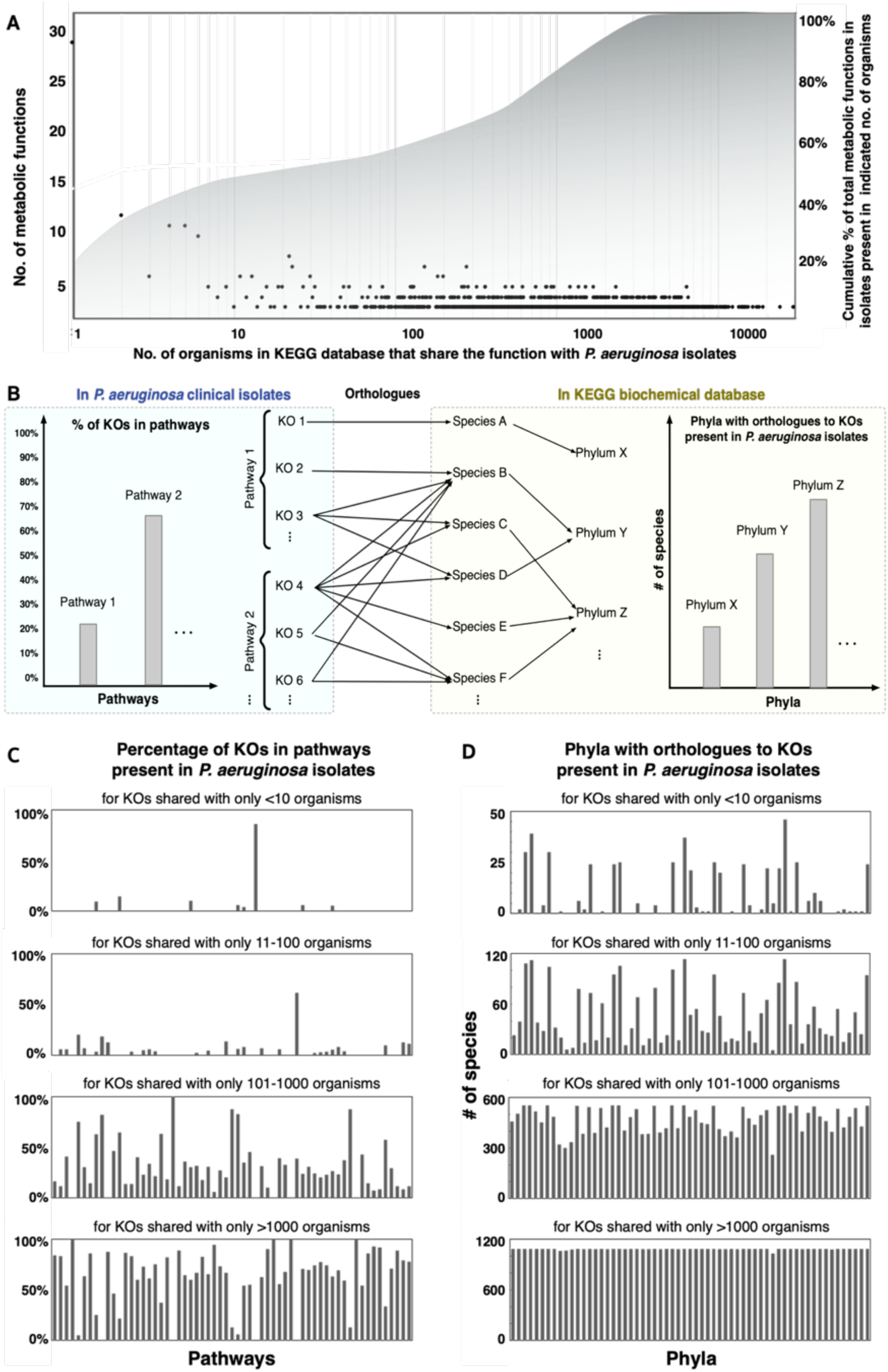
Functional annotations shared between the clinical *P. aeruginosa* isolates and other organisms. **(A)** Distribution of shared KEGG orthologues (KO) based on the number of non-pseudomonas organisms that contain the same KO. The area plot shows the cumulative number of functions that increase with increasing number of organisms. **(B)** A schematic of the search process for shared KOs with other organisms in the KEGG database. **(C)** Distribution of shared metabolic functions across different pathways. Only the pathways that shared >50% of the total KOs in that pathway with other organisms are displayed. With the increasing generality (# of organisms), increasing numbers of pathways are shared between the isolates and other organisms. **(D)** Distribution of shared metabolic functions across different phyla. With the increasing generality (# of organisms shared), the distribution of shared KOs tends to converge to a more uniform distribution across different phyla.

We calculated the fraction of KOs associated with a KEGG pathway that is present in the *P. aeruginosa* isolates as well as the number of species in a given phylum that contains orthologues to a given KO present in the isolates (Figures 2B-D). We calculated the fraction of KOs in a given pathway that is shared between the *P. aeruginosa* isolates and other organisms as a function of the number of organisms with shared functions. The number and diversity of the shared pathways increase with increasing number of organisms, but they uniformly belong to all the pathways (see Figure 3C). When we binned the KOs shared with increasing numbers of species (10-100, 100-1000, and more than 1000), we observed that the diversity of the non-*Pseudomonads* to which the isolates share their metabolic functions increased, but distribution across different phylogenetic lineages also becomes more uniform (see Figure 3D). This analysis helps to identify metabolic pathways that are unique to the *Pseudomonas* isolates we evaluated as well as which metabolic pathways are shared across multiple phyla.

Some of the major pathways that the clinical isolates share with less than 100 other species are Novobiocin biosynthesis and Phenazine biosynthesis. Novobiocin is a dibasic acid, which, like other aminocoumarin antibiotics, inhibits bacterial DNA synthesis by targeting the bacteria DNA gyrase and the related enzyme DNA topoisomerase IV. Although this antibiotic was effectively used to treat infections by Gram-positive bacteria like many Staphylococci ^33,34^, novobiocin has limited activity against Gram-negative organisms (like *P. aeruginosa*). This difference is due to the presence of the lipopolysaccharide-containing outer membrane that acts as a permeability barrier. *P. aeruginosa* has previously been observed to be resistant to novobiocin, mainly attributed to its MexAB-OprM multidrug efflux system that acts on this antibiotic ^35^. Other studies showed that novobiocin is responsible for binding to and inactivating the *nalD* gene that can repress the efflux pump, and therefore, contributes to intrinsic multidrug resistance in *P. aeruginosa* ^36^.

*P. aeruginosa* uses many secondary metabolites that act as virulence factors and negatively affect prokaryotic competitors and eukaryotic hosts through growth inhibition or cell death ^37^. Phenazines, such as pyocyanin, are redox-active, colored heterocyclic compounds and are responsible for the green fluorescence of *P. aeruginosa*. They play important roles in electron cycling, oxidative stress, and iron acquisition ^38,39^. Phenazine biosynthesis is responsible for promoting antibiotic tolerance and toxin production in *P. aeruginosa*. It also enhances the fitness of *Pseudomonas* in a biofilm environment and controls the production of many virulence factors^40,41^.

While the isolates shared most of their metabolic functions with many other non-*Pseudomonas* organisms, we observed that the annotated isolate genomes introduce several unique metabolic functions (KEGG orthologues) into the currently annotated *P. aeruginosa* pan-genomic landscape. Figure 4A shows the increasing addition of metabolic functions to the *P. aeruginosa* pan-genome with the increasing number of isolates sampled at random. These data have a large distribution because the number of combinations from which one can choose *n* number of isolates out of 25 is large, and each isolate contributes varying numbers of unique reactions toward the sample combinations. In total, the 25 clinical isolates introduce 66 metabolic functions, each of which is unique to a single clinical isolate. These KOs belong to aminobenzoate degradation; C5-Branched dibasic acid metabolism; Caffeine metabolism; Chlorocyclohexane and chlorobenzene degradation; Fructose and mannose metabolism; Glycine, serine and threonine metabolism; Inositol phosphate metabolism; Lysine biosynthesis; Phenylalanine, tyrosine and tryptophan biosynthesis; Purine metabolism; Pyruvate metabolism; Tryptophan metabolism; and Tyrosine metabolism, prominently. These KOs are not currently present in the annotated genome of *P. aeruginosa* in the KEGG database and were matched through sequence similarity against other organisms, mostly from Actinobacteria, Cyanobacteria, Euryarchaeota, Firmicutes, and Proteobacteria phyla (see Supplemental Information 2 for a complete list). While most previously characterized *Pseudomonas* strains do have some genes associated with many of these biosynthetic pathways, the newly identified genes in the clinical isolates perform additional or complimentary functions in these pathways. Since *P. aeruginosa* has extensive accessory genomic content within its pangenome, it is to be noted that the pangenome size will keep increasing with each additional genome sequenced and annotated over time. This analysis gives additional insight into the possibilities of finding novel metabolic functionalities in new strains isolated from previously unexplored sources.

**Figure 4:**
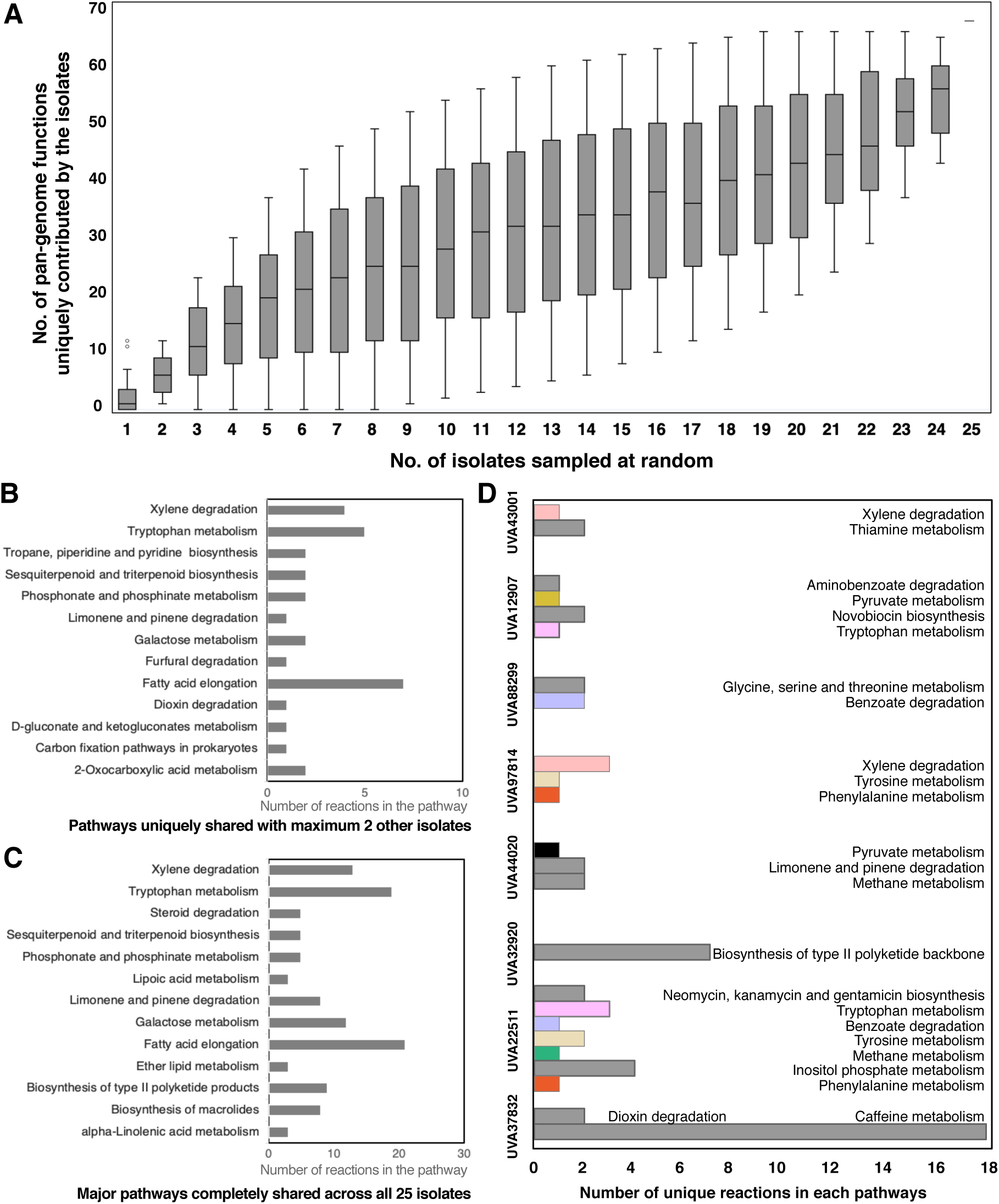
Analyses of the functional content of the *P. aeruginosa* clinical isolates. **(A)** Box-plot illustrating the distribution of unique metabolic functions (KEGG orthologues) contributed by a specific number of isolates that are selected at random. (**B** and **C**) Comparison of consensus and unique metabolic pathways across *P. aeruginosa* isolate models. (**B**) Pathways that are uniquely shared with less than two other isolates. The numbers below each pathway name correspond to the number of reactions in the pathway that are uniquely shared. (**C**) Major pathways that are completely shared by all the 25 clinical *P. aeruginosa* isolates. The numbers below each pathway name corresponds to the number of reactions in the pathway that are shared. (**D**) Unique metabolic functions in each of the clinical *P. aeruginosa* isolates. Pathways colored in grey are present in only one isolate. Pathways shared with at least one other isolate are colored.

Each of the isolates shares metabolic functions with very few of the other 24 isolates (one or two) uniquely. Figure 4B shows the distribution of reactions in different pathways that the isolates uniquely share with only one or two other isolates. Among these pathways, Fatty acid elongation, Xylene degradation, and Tryptophan metabolism contain the highest number of reactions.

In contrast, a total of 34 pathways are shared across all the clinical isolates. There are 13 metabolic pathways for which more than 20% of the reactions in that pathway are shared across all 25 isolates (Figure 4C). These include a total of 114 reactions in Xylene degradation, Fatty acid elongation, Galactose metabolism, and Tryptophan metabolism, among others. A list of the unique and shared reactions and the pathways these reactions belong to are presented in Supplementary Information 3.

The unique and common shared metabolism between clinical *P. aeruginosa* isolates has significant importance in guiding any future therapeutic development. While new drug design or repositioning strategies should be efficient enough to target unique metabolic traits of the clinically relevant isolates, their mechanism of action should also target a diverse set of strains to be effective as a treatment. The presence of unique functional traits in the metabolism of individual isolates creates a challenge for any therapeutic development effort. Therefore, it is important to characterize the metabolic functions unique to each of the isolates. Figure 4D shows the pathways in which unique reactions appear in the isolate genome annotations. Some of the isolates do not have any unique reactions, but eight of the isolates contain unique reactions in several pathways that are not shared by any other isolate. Interesting observations include the Biosynthesis of type-II polyketide backbone in UVA32920, and caffeine metabolism in UVA37832, among others. Bacterial aromatic polyketides, with their diverse structure, are involved in diverse biological activities, including producing antimicrobial component and deterrent molecules to outcompete other organisms, host immunosuppression, and virulence ^42–46^. Caffeine is known to inhibit the capability of *P. aeruginosa* to synthesize virulence factors and form biofilm by affecting the swarming motility and quorum sensing ^47–49^. Degradation of caffeine is known to be present in several *Pseudomonas* species ^50,51^, but the completeness of the caffeine metabolism pathway in only one of the isolates (UVA37832) potentially indicates its robustness against caffeine’s inhibitory action fitness and virulence in a clinical setting. Many of the unique pathways in other isolates are involved in extracellular signaling, regulation, as well as quorum sensing.

### Genome-scale metabolic network reconstructions of the isolates

To quantify the functional impact of these differences in metabolic gene content, we generated genome-scale metabolic network reconstructions for each of the clinical isolates. Metabolic network reconstructions combined with constraint-based analyses allow for a quantitative exploration of the functional repertoire and diversity of biological systems ^52–54^. We used the previously published genome-scale metabolic network reconstruction of *P. aeruginosa* PA14, iPau21 ^55^, as the backbone on which the additionally annotated metabolic reactions were added to generate the draft reconstructions of the isolates. The *P. aeruginosa* PA14 metabolic model was also amended with additional annotated reactions which were absent in iPau21. Each of the models was checked for reaction mass balance. The total number of reactions, unique reactions, genes, and metabolites are shown in Table 1. The Systems Biology Markup Language (sbml) version of all the reconstructions are available at https://anonymous.4open.science/r/PA_clinical_isolate_reconstructions/.

**Table 1:**
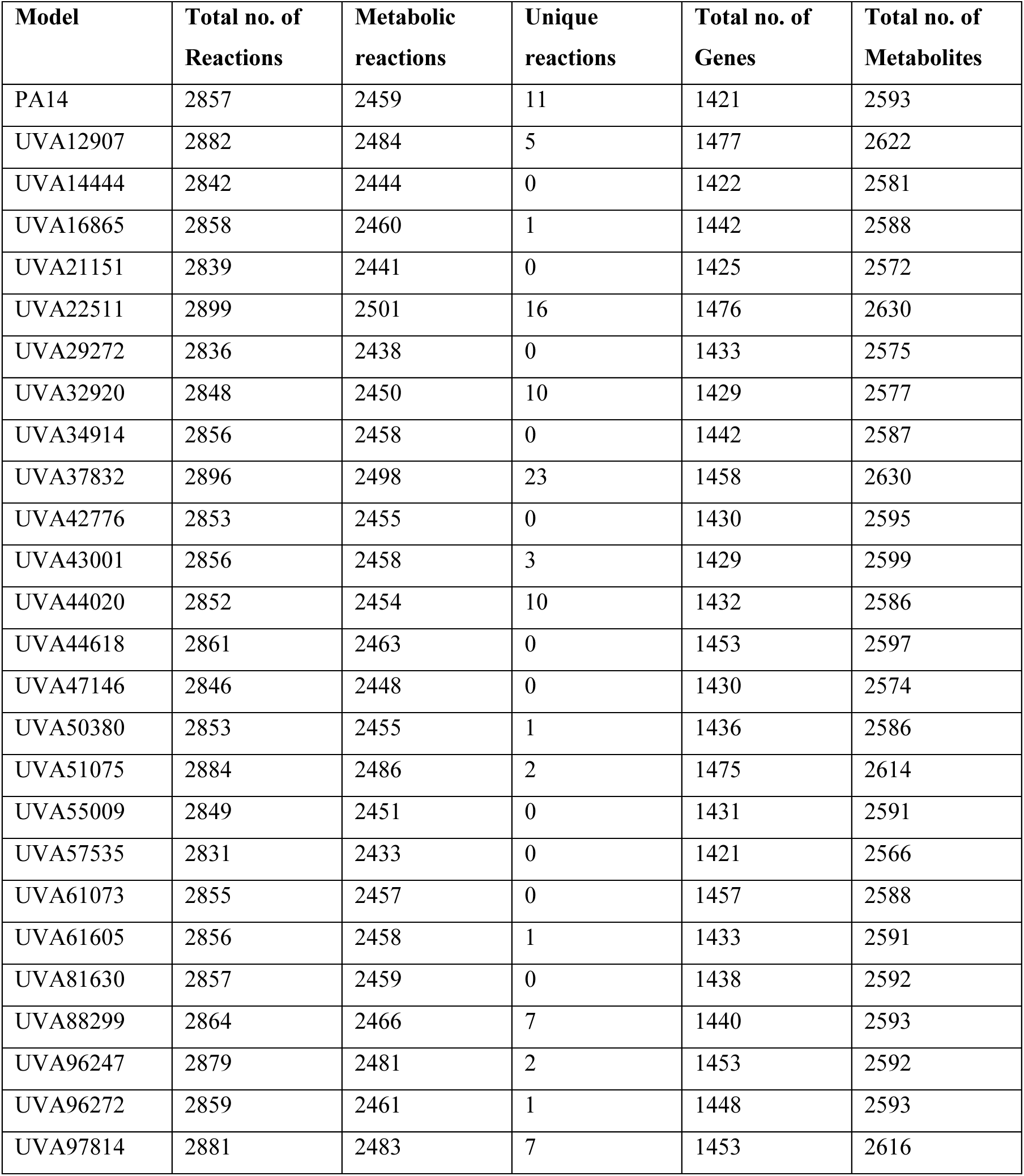
Properties of the metabolic reconstructions of the *P. aeruginosa* clinical isolates.

### Diversity of metabolic flux states of the isolate metabolic reconstructions

The isolate metabolic network reconstructions were simulated in *in silico* Synthetic Cystic Fibrosis Media (SCFM) as described previously ^55^ to generate 500 flux sample predictions for each model. SCFM is a physiologically relevant medium that elicits more realistic metabolic behavior of the isolates in an infection setting. To compare the flux sampling results across isolates, a non-metric multidimensional scaling (NMDS) method was used. Supplementary Figure 1 shows the flux sampling results in a 2-dimensional plane. Here, each data point represents a functional metabolic snapshot of the flux sampling data. The data points are colored for each of the isolates, and we observe that while most of the isolates as well as the reference PA14 model overlap in certain functional metabolic states, each of them has a distinct set of flux samples, as shown by the outward rays from the center. To ensure convergence of the flux sampling algorithm, we also enumerated 10,000 flux samples for each of the isolates and performed non-metric multidimensional scaling (shown in Supplementary Figure 2). The NMDS distances and the correlation with phenotypic data do not change significantly when a higher number of flux samples are performed.

To evaluate if the flux sampling results correlate with the isolate phenotypes and patient metadata, the NMDS plot was color-coded with each of the phenotypic categories in Figure 5. Some phenotypic traits correlate with the flux samples better than others. For example, the correlation values of the isolate collection site (0.46) and patients’ sex (0.44) are noticeably higher than the correlation to other traits like comorbidities and isolate morphology. While these calculated correlation values are possibly confounded by many unknown factors, the higher correlation values between isolates’ functional landscape and patient demography and isolation site indicate that specific host environment may be a critical factor in shaping *P. aeruginosa* metabolism. Different environmental niches within the host have been observed to rewire the metabolism and pathogenicity of *Pseudomonas* strains in recent studies ^56–59^. The kind of modulation of metabolic functions in different host sites may have significant implications in the evolution of the pathogen within an extended timeframe of infection, which has also been observed in laboratory studies ^60^.

**Figure 5:**
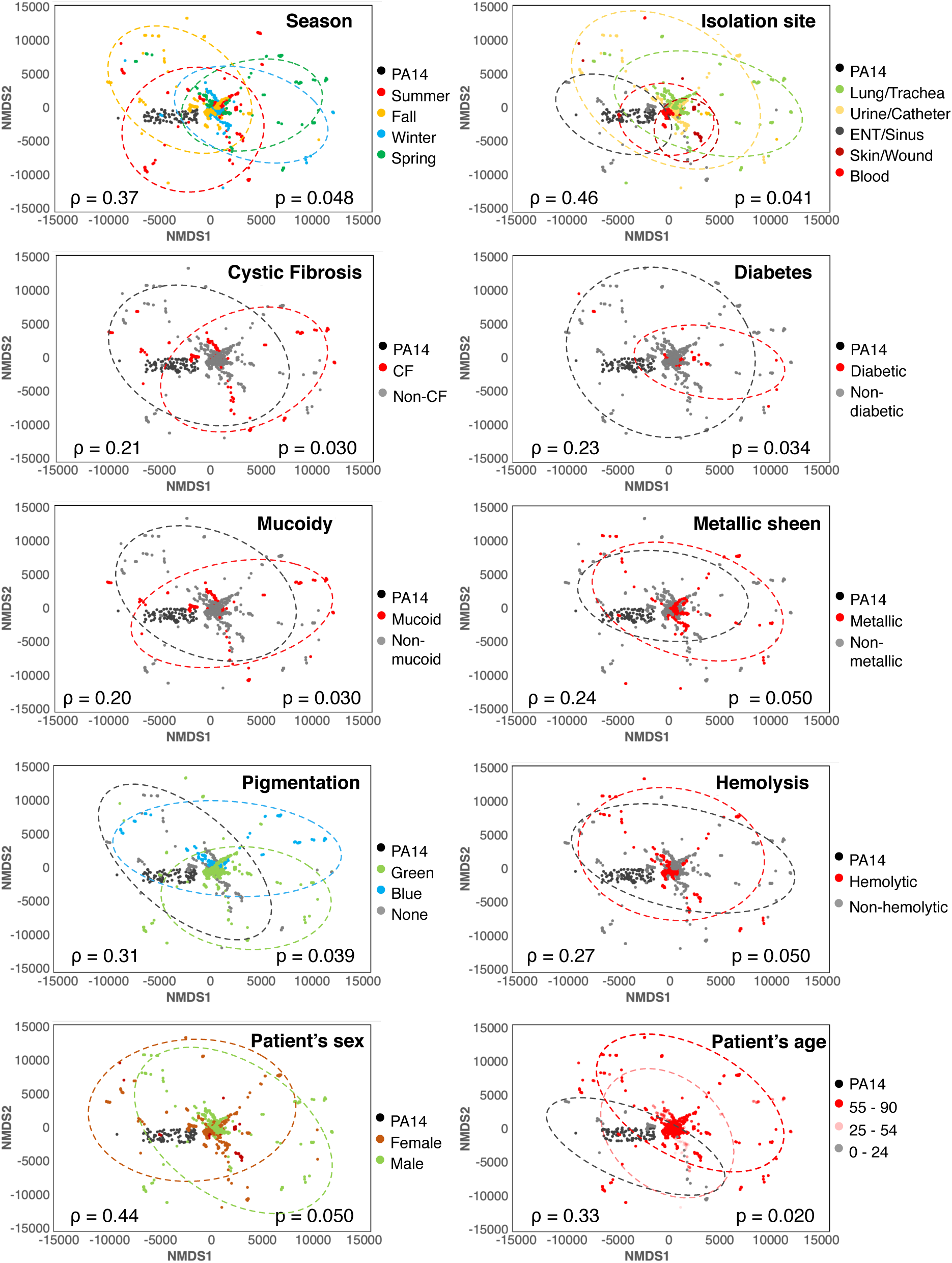
NMDS plots of the flux sampling results from the isolates’ metabolic network reconstructions color-coded according to isolate collection sites and season, patient demographic profile and isolate morphology.

## Conclusions

We performed a multi-faceted analysis of a representative group of *P. aeruginosa* clinical isolates by employing whole genome sequencing, phenotypic and genotypic clustering, functional annotation and analyses on core, accessory, and unique traits in the *P. aeruginosa* pan-genome, and genome-scale metabolic network modeling. This study demonstrates the importance of an in-depth study of isolate sources, patient metadata, and morphological phenotypes and their connections to the diverse metabolic landscape of clinical *P. aeruginosa* isolates. Through our multi-dimensional approach, we have characterized the diverse genotypes and metabolic functions across the clinical isolates with a wide range of phenotypes. We utilized a publicly available high-quality clinical data set and developed several analysis pipelines that can be employed for other human pathogens.

The extensive work by Dunphy *et al* ^10^ to assemble the collection of 971 clinical isolates from the UVA Health System laid the groundwork for the current study. To select a representative group of isolates from the huge collection, the use of stratified random sampling allowed us to obtain a sample population of isolates that best represents the phenotypic diversity of the entire isolate collection. Whole genome sequencing allowed us to compare the genomic content and therefore differentiate the selected isolates based on sequence similarity (Figure 1). We found that the genomic differences among the set of 25 isolates were lower than that observed in other *P. aeruginosa* pangenomic studies ^13,19,20,61^. The allelic profiles of the sequenced isolates were not too diverse from each other, as observed in Figure 2. This result is not unexpected considering this study involved isolates from one single geographical location. Also, a strong correlation between the genomic content, multi-locus sequence type, and phenotypic characteristics was not observed in our study. For a more complete picture of their functional diversity and uniqueness, annotation of the genomic sequences was important. It revealed not only how the isolates are similar or unique compared to each other as well as the reference *P. aeruginosa* PA14 strain, but also how the clinical isolates expand the entire metabolic landscape of the *P. aeruginosa* pan-genome, sharing metabolic functions from a diverse range of species (Figures 3 and 4). With future sequencing and further annotation of the many *Pseudomonas* strains, certainly, these metabolic functions will certainly be identified in other isolates.

Genome-scale metabolic models are powerful platforms for analyzing the active metabolism of a species by using computational tools based on constraint-based modeling methods ^52,53,62–65^. In this work, we reconstructed the metabolic networks of each of the isolates from the genome annotation and built on top of a highly curated, genome-scale metabolic model of the PA14 strain, *i*Pau21 ^55^. While we recognize that there are many non-metabolic functional differences that might be present among the different clinical isolates, we primarily focus on metabolic reactions that distinguish their behavior. We enumerated 500 flux samples for each of the isolate models to evaluate their relative distance based on a non-metric multidimensional scaling method. We observed that while most of the strains share much of their metabolic profiles, all of them show some unique metabolic functions (Figure 5). Correlation of the flux sampling results with phenotypic traits identifies patient sex and strain isolation site as the most important factors in shaping the pathogen metabolism. We also explored the distinguishing feature of the annotated KEGG pathways and performed the same non-metric multidimensional analysis based on the annotated functional content in each of the isolates (shown in Supplementary Figure 3). We observed that none of the isolate phenotypes or patient metadata correlate with the functional content-based dimensionalization significantly better than flux sampling data.

There are several areas where this study can be complemented by other *in vitro* analyses and high throughput analyses. For example, while the whole genome sequencing and annotation provide us with a more complete picture of the metabolic capabilities of the clinical isolates, a high-throughput transcriptomic study could reveal their metabolic adaptations in an infection setting when they are subject to a host immune system among other factors. Functional transcriptomics of the core and unique metabolism of the different isolates can also enable the evaluation of the strain-specific variations in virulence mechanism and adaptability. In addition, the semi-automatic curation of the isolate metabolic models can be further refined with high-quality gene essentiality data as well as substrate consumption and fitness profiles. Overall, this work paves the foundation for further integrative studies to understand the mechanisms and diversity of metabolic modulations in *P. aeruginosa* associated with the host environment.

## Methods

### Isolate collection

As described previously, a total of 971 clinical isolates of *P. aeruginosa* were collected from the UVA Health System Clinical Microbiology Laboratory between February 2019 and February 2020 ^10^. For each isolate, associated patient metadata (age, sex, isolation sites, and whether the patient had cystic fibrosis or diabetes mellitus), bacterial morphological phenotypes, and antimicrobial susceptibility profiles were recorded ^10^. Isolate sources that could not be classified as from the lung/trachea, urine/catheter, ENT/sinus, skin/wound, or blood were grouped as “other.” Supplementary information 4 lists all the isolates and their patient metadata and phenotypes used in this study.

### Stratified random sampling

The entire population of 971 isolates was divided into 10 homogeneous strata or subgroups according to the patient metadata (e.g., age, sex, isolation sites, presence of CF or diabetes mellitus, season) and isolate phenotypic properties (e.g., mucoidy, color, hemolytic property, and metallic sheen). Stratified random sampling ^66,67^ from each of the non-overlapping strata was performed. A Mixed-Integer Linear Programming (MILP) formulation was used to draw a random sample from the different strata separately.

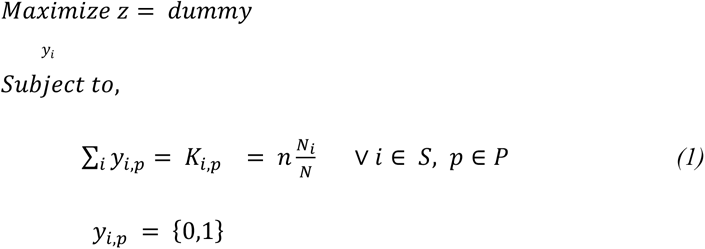

Where *y_i,p_* is the binary variable denoting the inclusion or exclusion of the isolate into the sampled set. S is the set of the strata for each phenotypic trait P. *K_i,p_* is the population proportion of the stratum *i* in the phenotype *p, n* is the total number of sample units available for allocation, and *n_i_* is the number of sample units to allocate to stratum *i*. This framework allows one to obtain an effect size from each stratum/phenotype separately and independently of other phenotypic properties. It also ensures that isolates are selected from every stratum instead of leaving out the minority sets that can happen in random sampling.

### Bacterial culture and genome sequencing

Selected *P. aeruginosa* isolates were transferred from frozen stocks into LB broth with shaking (37C, 150rpm, 16-24h). Cultures were plated on blood agar and cetrimide agar for phenotype verification. Cells were pelleted from cultures and DNA isolated using the DNeasy UltraClean Microbial Kit (Qiagen) according to manufacturer’s instructions. Purified DNA was quantified using the broad range dsDNA kit (DeNovix), libraries generated, and samples sequenced (2x151bp; 400Mbp/sample) on the Illumina NextSeq 2000 platform (SeqCenter; Pittsburgh, PA). Sequences are available at http://www.ncbi.nlm.nih.gov/bioproject/937715.

### Sequence analyses

FastQC v0.11.9 (bioinformatics.babraham.ac.uk/projects/fastqc/) was used to examine the quality of both the forward and reverse reads. Trim Galore v0.6.5 (bioinformatics.babraham.ac.uk/projects/trim_galore/) was used for automatic quality and adapter trimming of the sequences. Across all the paired isolate sequences, the median per base sequence quality score was above 32. For assembling the paired end reads to contigs, Velvet ^68^ was used with a max *k*-mer length of 101 bp. The resulting contigs were ordered against the reference sequence of *P. aeruginosa* PA14 genome (NCBI ID: NC_008463.1). Pairwise genome comparison was performed using ACT ^69^ after converting the multi-FASTA files generated by Mauve ^70^ to single-FASTA format files. Basic local alignment search (BLAST version 2.10.0) was used to create comparison files between the reference PA14 genome and the isolate genome sequences. BLAST Ring Image Generator (BRIG v0.95) ^29^ was used to generate the ring diagram showing the isolate whole-genome sequences in comparison with each other as well as the reference PA14 genome.

### Clustering and Correlation

Phylogenetic clustering was performed with MEGA11 ^27,71^ on a local workstation. The whole genome nucleotide sequence was translated to protein sequences before a BLASTp search (BLAST version 2.10.0) was performed across the isolates. The individual isolate protein sequence “.fasta” files were inputted to the MEGA alignment explorer and the MUSCLE ^72^ algorithm was used to align the sequences. Multiple Alignment Gap Opening penalty was set to 3 and the Multiple Alignment Gap Extension penalty was set to 1.8, according to recommended values ^28^. UPGMA Maximum Parsimony was used to evaluate the phylogenetic tree and an unrooted tree was generated^32^. For clustering of non-numeric categorical data like the phenotype and patient metadata corresponding to each of the isolates, a metric called “Category Utility” was used. Category Utility is a measure of “category goodness” that attempts to maximize both the probability that two objects in the same category have similar attribute values, and the probability that objects from different categories have different attribute values ^26^. The Category Utility value (CU) of a given clustering of a dataset is a numeric value that reflects how well the clustering is performed. Larger values of CU indicate better clustering.

Since data clustering is an NP-hard problem, there is no way to find an optimal clustering without examining every possible clustering. One way to increase the likelihood of an optimal clustering is to cluster the dataset multiple times with different initial cluster assignments. Then, the algorithm iteratively tries to find the best clustering of the data based on the initial cluster assignments. In this study, 125 different initial cluster assignments were attempted and the clustering with the highest CU value of 0.3999 was chosen as the optimal phenotypic clustering of the isolates.

Multi-Locus Sequence Typing (MLST) was evaluated for the isolate on the PubMLST database (https://pubmlst.org) with *Pseudomonas aeruginosa* as the genome assembly for the species ^30,73^. 19 of the 25 isolates showed a 100% identity match of the sequences for all the 7 housekeeping genes (*acsA, aroE, guaA, mutL, nuoD, ppsA,* and *trpE*) on the PubMLST database. For the six other isolates, the nearest match of sequence type was chosen (see Supplementary Information 1 for detailed results). Based on the MLST results, a Minimum spanning Tree (MST) was created with PHYLOViZ (online version as of August 2023) ^74^. Hierarchical clustering and entanglement scores were enumerated using the “dendextend” package^75^ version 1.17.1 in R version 4.1.2.

Non-metric multidimensional scaling (NMDS) ^76^ was used to reduce the dimensionality of the flux sampling results from each of the isolates. A Bray-Curtis distance metric ^77^ was used in the evaluation of the NMDS. A maximum iteration limit of 500 was set. The optimal stress value of the resulting NMDS was 0.1480.

### Functional annotation

The “.fasta” files containing the post-processed genome sequences were annotated based on the KEGG biochemical database (as of December 2021) ^31,32^. First, each of the sequence files was processed with Prodigal v2.6.3 ^78^ to translate to protein sequences and the coordinates for the coding sequences (CDS). A BLASTp search (BLAST version 2.10.0) across the entire KEGG prokaryotic database was performed for each of the protein-coding sequences using DIAMOND v2.0.14 ^79^. A percent identity >98%, e-value <1e-05, and a bit score >50 were used as thresholds for selecting best sequence similarity matches. Protein IDs from all the KEGG matches were then translated to KEGG Orthologue (KO) numbers after removing duplicates. KEGG reaction IDs were obtained by parsing the KO to reaction mapping.

### Genome-scale metabolic network reconstructions and analyses

A genome-scale metabolic network reconstruction was generated from the annotated genome sequence for each of the 25 isolates. The recently published genome-scale network reconstruction of *P. aeruginosa* PA14 (iPau21) by Payne et al. ^55^ was used as the foundation on which the isolate models were reconstructed. Since the base model contained reaction and metabolite IDs in the ModelSEED ^80^ namespace (as of March 2021), the annotated KEGG reactions and associated metabolite IDs were translated to ModelSEED IDs. The reactions that were annotated in each of the isolate genomes but were not present in the iPau21 model were added to the corresponding reconstruction. The models were checked for mass and charge imbalances. For generating flux samples, each isolate model was sampled with optGpSampler ^81^ algorithm in CobraPy v0.21.0 ^82^ for 500 times. The Jaccard distance ^83^ of phenotypic categories of the isolates was calculated in a pairwise fashion. The NMDS distance was calculated as the distance between the median NMDS coordinates of isolate pairs. Spearman’s correlation ^84^ was used to calculate a *p*-value for the relationship between NMDS distance between isolate pairs and their phenotypic distance (Jaccard index).

## Data availability

The datasets generated and analyzed during the current study are available at http://www.ncbi.nlm.nih.gov/bioproject/937715. The Systems Biology Markup Language (sbml) version of all the genome-scale metabolic network reconstructions of clinical isolates are available at https://anonymous.4open.science/r/PA_clinical_isolate_reconstructions/.

## Funding

We acknowledge support by the National Institutes of Health (grant number R01-AI154242).

## Author contribution

MMI: Experimental design, Data collection, Data analysis, Software, Original Draft, Writing, Review and Editing; GLK: Experimental design, Data collection, Writing, Review and Editing; EMG: Writing, Review and Editing; JBG: Project administration, Writing, Review and Editing; JAP: Project administration, Writing, Review and Editing.

## Notes

### Competing Interest Statement

The authors have declared no competing interest.

