## Supplementary figures and images for "Model-driven characterization of functional diversity of *Pseudomonas aeruginosa* clinical isolates with broadly representative phenotypes"

### Supp_figure_ S1.tiff

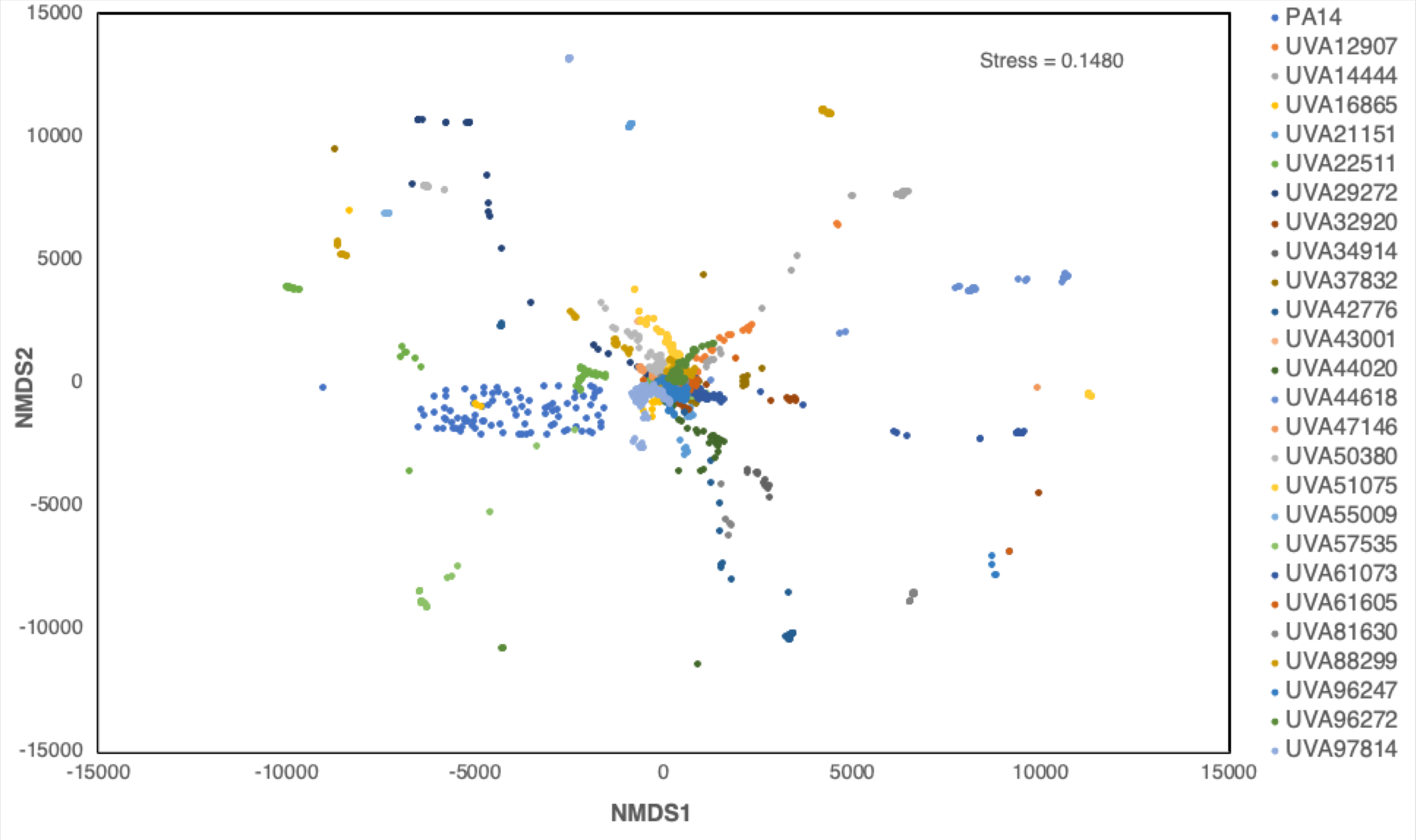

### suppl_fig_1.tiff

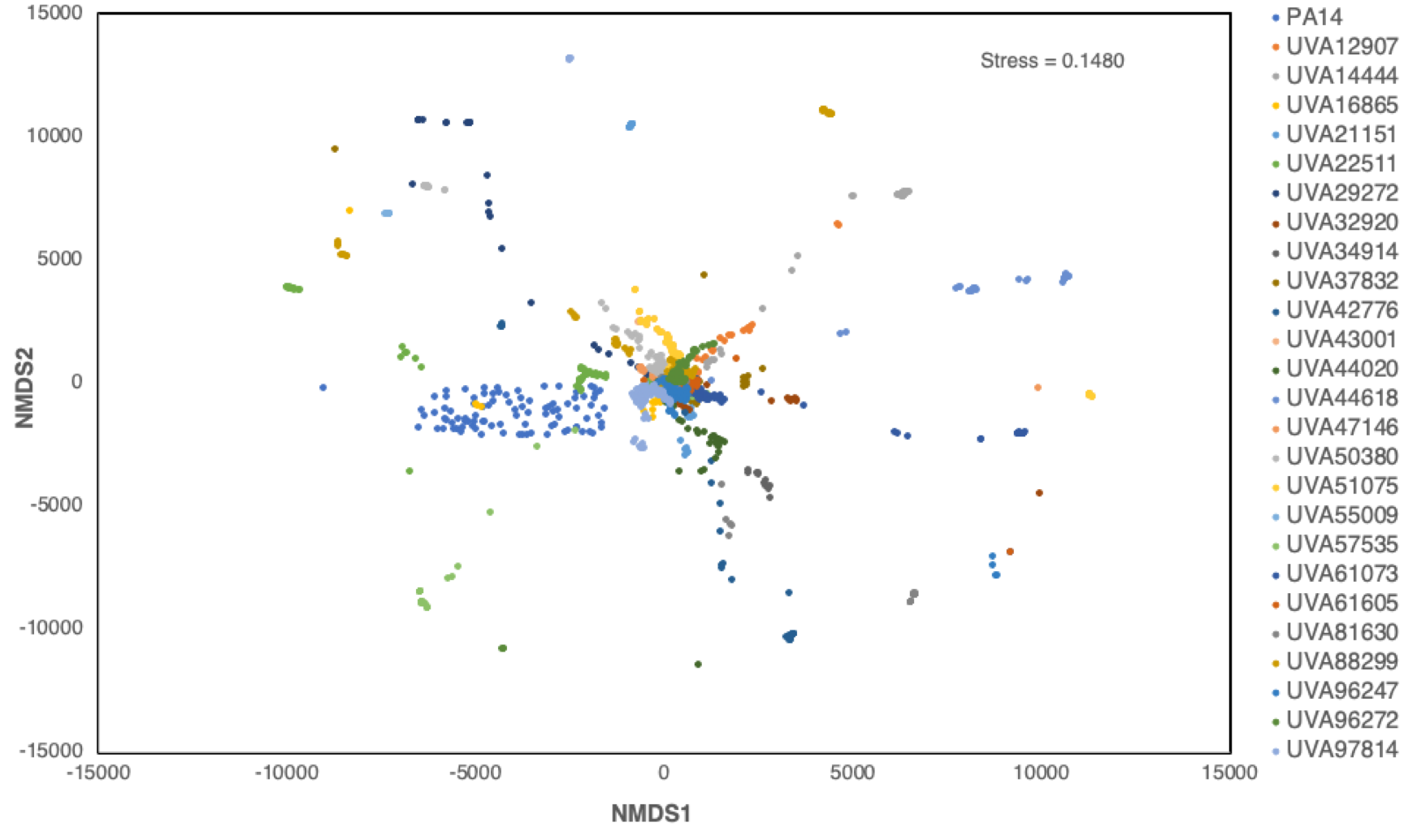

### suppl_fig_3.tiff

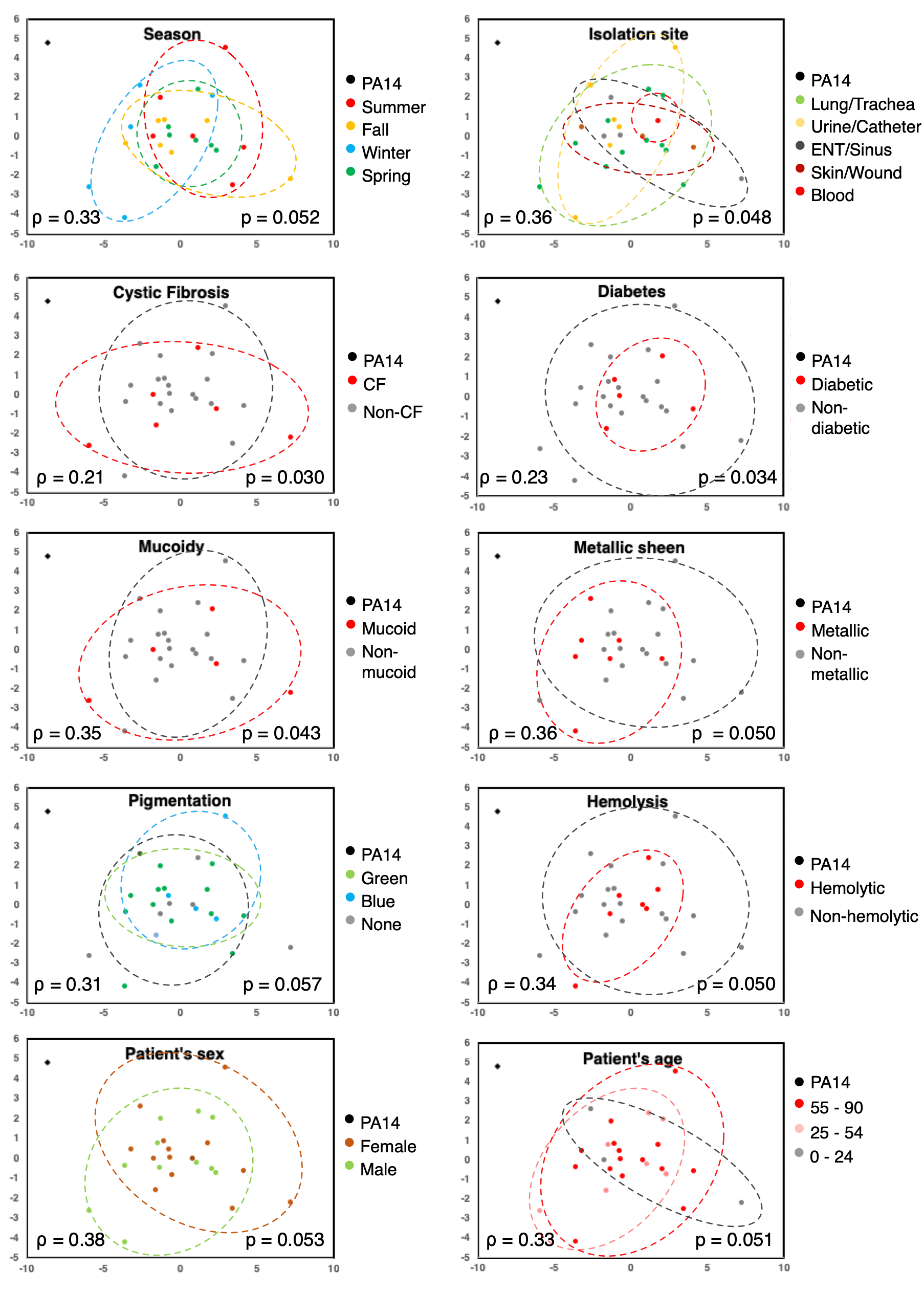

### supple_fig_2.png

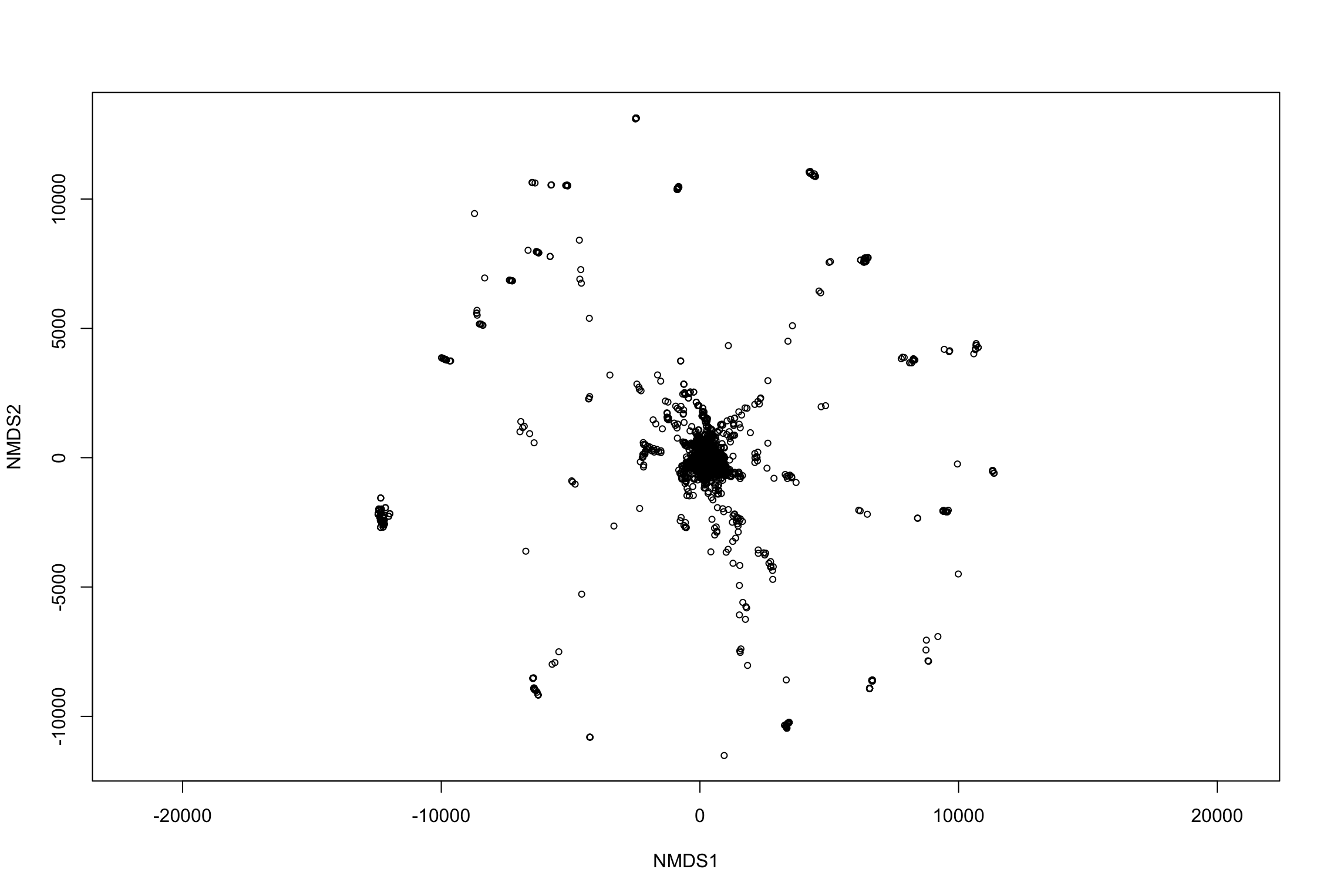
